# Neonatal Subarachnoid Hemorrhage Disrupts Multiple Aspects of Cerebellar Development

**DOI:** 10.1101/2023.02.10.528048

**Authors:** David F. Butler, Jonathan Skibo, Christopher M. Traudt, Kathleen J. Millen

## Abstract

Over the past decade, survival rates for extremely low gestational age neonates (ELGANs; <28 weeks gestation) has markedly improved. Unfortunately, a significant proportion of ELGANs will suffer from neurodevelopmental dysfunction. Cerebellar hemorrhagic injury (CHI) has been increasingly recognized in the ELGANs population and may contribute to neurologic dysfunction; however, the underlying mechanisms are poorly understood. To address this gap in knowledge, we developed a novel model of early isolated posterior fossa subarachnoid hemorrhage (SAH) in neonatal mice and investigated both acute and long-term effects. Following SAH on postnatal day 6 (P6), we found significant decreased levels of proliferation with the external granular layer (EGL), thinning of the EGL, decreased Purkinje cell (PC) density, and increased Bergmann glial (BG) fiber crossings at P8. At P42, CHI resulted in decreased PC density, decreased molecular layer interneuron (MLI) density, and increased BG fiber crossings. Results from both Rotarod and inverted screen assays did not demonstrate significant effects on motor strength or learning at P35-38. Treatment with the anti-inflammatory drug Ketoprofen did not significantly alter our findings after CHI, suggesting that treatment of neuro-inflammation does not provide significant neuroprotection post CHI. Further studies are required to fully elucidate the mechanisms through which CHI disrupts cerebellar developmental programming in order to develop therapeutic strategies for neuroprotection in ELGANs.

## Introduction

Despite improvements in survival rates for extremely low gestational age neonates (ELGANs; <28 weeks gestation), approximately 50% of survivors go on to have significant neurodevelopmental dysfunction including cognitive, learning, and/or motor deficits. Many of these deficits do not seem to be explained by cerebral white matter injury and/or IVH, suggesting an alternate mechanism of injury (Nosarti et al., 2007; Limperopoulos et al., 2014). Recent data suggests that injury to the cerebellum contributes to neurodevelopmental deficits and that developmental disruption of the cerebellum has widespread effects on neurologic function (Haines et al., 2013; Dijkshoorn et al., 2020; Brossard-Racine and Limperopoulos, 2021a; Spoto et al., 2021).

Between 20 and 40 weeks of gestation, cerebellar growth is vigorous, with the volume of the cerebellum increasing at least fivefold. This rapid growth is almost exclusively driven by massive proliferation granule neuron progenitors in the external granule layer (EGL). These progenitors will then differentiate into cerebellar granule cells (GC) and migrate into to their final location beneath the developing Purkinje cells via fibers of Bergman Glial cells to form the internal granule cell layer (IGL) (Rakic, 1971; Abraham et al., 2001; Volpe, 2009; Consalez et al., 2020). As the cerebellum rapidly grows, the developing cerebellar circuitry is also undergoing rapid remodeling and maturation. Disruptions of this early circuitry alter long term cerebellar circuit maturity (Haldipur et al., 2011; Barron and Kim, 2020; van der Heijden et al., 2021; van der Heijden and Sillitoe, 2021) and deranged cerebro-cerebellar neuronal circuitry causing impaired cerebral development and profound neurologic dysfunction (Limperopoulos et al., 2010; Limperopoulos et al., 2014; Wang et al., 2014; D’Mello and Stoodley, 2015; Stoodley and Limperopoulos, 2016; Brossard-Racine and Limperopoulos, 2021a).

Injury to the neonatal cerebellum has been increasingly recognized in the ELGANs population. Many factors, including nutritional deficits, infection, inflammation and glucocorticoid exposure can disrupt cerebellar development in preterm infants (Buddington et al., 2018; Iskusnykh et al., 2018; Gano and Barkovich, 2019; Chizhikov et al., 2020; Iskusnykh et al., 2021; Iskusnykh and Chizhikov, 2022). However, cerebellar hemorrhagic injury (CHI) is a relatively frequent finding in preterm infants are more even more common in severely premature infants (Steggerda et al., 2009; Boswinkel et al., 2019; Villamor-Martinez et al., 2019). Radiologic evidence of cerebellar subarachnoid and parenchymal hemorrhages correlates with poor neurodevelopmental outcomes (Johnsen et al., 2005; Steggerda et al., 2009; Zayek et al., 2012; Matsufuji et al., 2017; Boswinkel et al., 2019; Brossard-Racine and Limperopoulos, 2021b). Hemorrhage is also highly associated with cerebellar hypoplasia (Volpe, 2009; Aldinger et al., 2019; Gano and Barkovich, 2019; Scelsa et al., 2022). Additionally, documented fetal cerebellar hemorrhage has been associated with cerebellar hypoplasia and neurodevelopmental deficits in full term children (Poretti et al., 2008). Furthermore, cerebellar hypoplasia on MRI has been shown to more predictive of neurodevelopmental deficits than supratentorial injuries such as intraventricular hemorrhage or white matter injury (Limperopoulos et al., 2007).

While hemorrhage has been associated with cerebellar hypoplasia, the underlying injury mechanism is not fully understood. Extravasated blood and/or blood components have been shown to be injurious to multiple cell populations in the developing cerebral cortex, including neurons, glia, and oligodendrocytes (Xue et al., 2003; Vinukonda et al., 2012). Lypophosphatidic acid (LPA), a component of serum, induces hydrocephalus when injected into the lateral ventricles of embryonic mice due to ependymal injury and progenitor delamination into the lateral ventricles (Yung et al., 2011). Additionally, hemosiderin deposition may contribute to increases in reactive oxygen species (Volpe, 2009). However, the cerebellar-specific literature is very limited. We previously reported that a rabbit model of cerebellar subarachnoid hemorrhage induced in neonatal rabbits via systemic glycerol administration is invalid since glycerol itself is toxic to cerebellar development even in the absence of hemorrhage (Traudt et al., 2014). One of the first mouse models of neonatal cerebellar hemorrhage conducting intraventricular injection of bacterial collagenase into the fourth ventricle of early post-natal mice, when maximal EGL proliferation occurs in this species (Yoo et al., 2014).

Subsequent hematomas in the fourth ventricle and on the cerebellar surface caused decreased granule cell density, decreased cerebellar volume and resulted abnormal open field testing behavior, and motor impairment on Rotarod testing in older mice (Yoo et al., 2014), which is exacerbated by inflammation (Tremblay et al., 2017). The effects of isolated posterior fossa blood exposure in the absence of intraventricular hemorrhage on cerebellar development have never been directly tested *in vivo*.

In this study, we developed a novel model of isolated posterior fossa subarachnoid hemorrhage in neonatal mice to directly examine the effects of blood exposure on cerebellar development. Our model avoids direct injury to the cerebellar parenchyma and isolates injury related to blood components. We examined acute and long-term effects on cerebellar cellular biology, anatomy, and animal behavior following cerebellar. We hypothesized that exposure of the developing cerebellum to blood would result in disruption to multiple developmental programs leading to long-term cellular/anatomic abnormalities and motor deficits. Secondarily, we hypothesized that neuroinflammation may strongly contribute to the underlying injury mechanism and that treatment with an anti-inflammatory may be neuroprotective.

## Materials and Methods

### Animals

This study was carried out in accordance with the recommendations in the Guide for the Care and Use of Laboratory Animals of the National Institutes of Health and was approved by the Institutional Animal Care and Use Committee (IACUC) at University of Washington (Protocol Number: IACUC00006). Mice from the CD1 strain were bred and housed in Optimice cages with aspen bedding at the Seattle Children’s Research Institute specific pathogen-free (SPF) vivarium facility on a 14/10 hour light/dark cycle. All mice were monitored closely by both laboratory and veterinary staff. The day of birth was designated as postnatal day 0 (P0).

### Posterior fossa injection procedure

On P6, mice underwent posterior fossa injections under anesthesia with isoflurane. A single 8-10mm vertical incision was made overlying the posterior calvaria (**Figure 1A**). Trunk blood from an adult CD1 mouse was obtained and anticoagulated with heparin at a ratio of 100µl heparin to 900µl of blood. For the CHI group, 20µl of trunk blood was injected into the posterior fossa with a 28-gauge insulin needle at the midline position, 2.5mm rostral from Lambda under direct visualization. Control group animals underwent posterior fossa injection with 20µl of heparin containing artificial cerebrospinal fluid (aCSF) (Wei and Ramirez, 2019) as outlined above.

**Figure 1.**
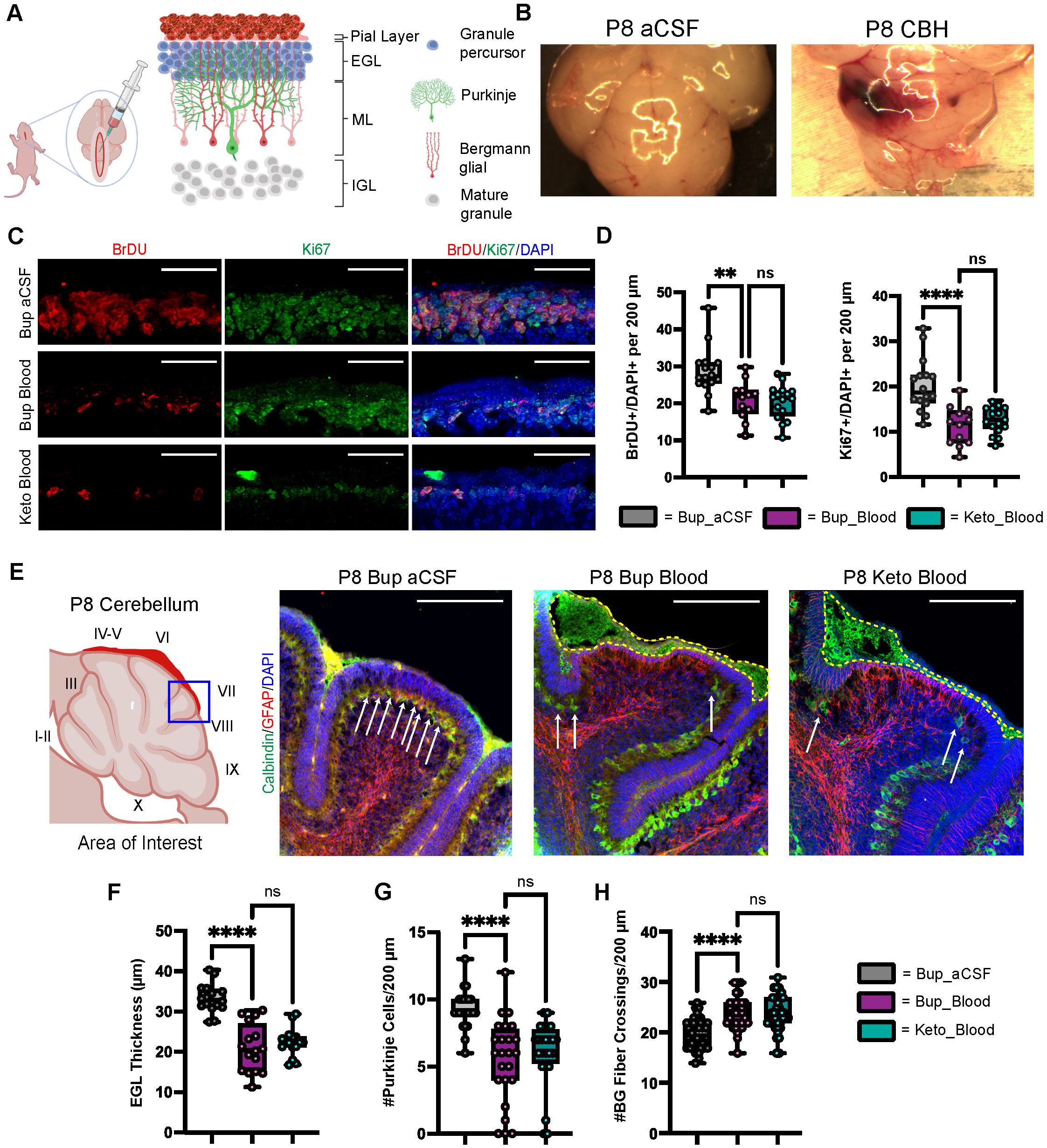
CHI results in decreased BrDU and Ki67 labeling indices, thinning of the EGL, decreased PC density, and increased BG fiber crossings at P8. **(A)** Schematic illustrating isolated posterior fossa hemorrhage model with typical P6 cerebellar architecture. **(B)** Representative comparative images of gross findings between aCSF and CHI groups; note large hematoma formation overlying left cerebellar hemisphere in the CHI specimen. **(C)** Representative confocal images at 40X of BrDU and Ki67 staining within the EGL. Only cells within the EGL were included in analysis. Scale bars are 25 *μ*m. **(D)** Quantification of labeling indices for BrDU (left) and Ki67 (right); data obtained from 200 µm segment from lobule VI and VII (N = 2, n = 12 for Bup_Blood; N = 3, n = 16 for Bup_aCSF and Keto_Blood). **(E)** Representative confocal images at 20X magnification of lobule VII demonstrating differences in local findings including presence of large RBC collections (yellow dashed outline), PC density (white arrows), thickness of EGL, and architecture of BG fibers. Scale bars are 100 *μ*m. **(F)** Quantification of EGL thickness for all treatment groups (N = 4, n = 16). **(G)** Quantification of PC density in lobule VI and VII for all treatment groups (N = 3, n = 24). **(H)** Quantification of BG fiber crossings in lobule I, VII-IX for all treatment groups (N = 3, n = 27). Significance defined as p < 0.05 by one-way ANOVA with Tukey post-test. Data are presented as SEM with minimum and maximum values. N = number of animals per group included for analysis, n = number of section segments per group included for analysis.

Following injection, the needle was gently removed, and the skull defect was covered with Vetbond(tm) tissue adhesive (3M). Skin closure was performed with 7.0 Ethilon® nylon suture (Ethicon). Animals received either buprenorphine (2µg/g of body weight) or ketoprofen (5µg/g of body weight) for analgesia. Mice were warmed and monitored closely during the entire procedure and recovered under a heat lamp until they were noted to have consistent spontaneous movements prior to return to their home cages.

### Sample preparation

On P8 or P42, mice were pulsed with 5-bromo-2’-deoxyuridine (BrdU) via intraperitoneal injections (10µg/g of body weight) 2 hours prior to sacrifice. On P8, mice were sacrificed via rapid decapitation, tissue was harvested and then fixed in 4% paraformaldehyde (PFA) overnight. On P42, mice underwent both cardiac fixation with 4% PFA infusion and drop fixation in 4% PFA overnight. At both timepoints, tissue was then equilibrated in 30% sucrose made in phosphate-buffered saline (PBS). Cerebellar tissue was isolated, placed in Tissue-Tek® O.C.T compound and stored at -80 °C. Sections were performed at 18-20µm on a freezing microtome.

### Immunohistochemistry (IHC)

Sections were dried at room temperature for 60 minutes, baked at 45 °C for 30 minutes, and washed in PBS. Sections undergoing BrdU staining for were bathed in 2N HCl for 30 minutes. All sections underwent antigen retrieval by boiling sections in 10mM sodium citrate for 10-15 minutes. Sections were then washed three times with PBS with 0.125% Triton X-100 (PBX) and blocked in 5% donkey serum in PBX in a humidified chamber for 90 minutes at room temperature. Sections were incubated at 4 °C with primary antibodies in a humidified chamber overnight. The following day, sections were washed in PBX and then incubated with species-specific secondary antibodies conjugated with Alexa 488, 568 or 594 fluorophores (Invitrogen) for 90 minutes at room temperature. Sections were counterstained with DAPI (4’,6-Diamidino-2-Phenylindole, Dihydrochloride; Invitrogen; D1306) and cover-slipped with Fluoromount-G (Southern Biotech, Cat. No. 0100-01). Primary antibodies used included: rat anti-BrdU (1:100, Abcam), mouse anti-Calbindin (1:20, Abcam), rabbit anti-GFAP (1:1500, Dako), rabbit anti-Ki67 (1:150, Novus), mouse anti-Parvalbumin (1:5000, Swant), rabbit anti-BLBP (1:400, Abcam).

### Rotarod assay

At P35, all mice underwent motor learning and coordination testing using the Rota-Rod 2 (Med Associates Inc.). On day 1 of testing, each mouse underwent three trials, and on days 2 and 3, each mouse underwent two trials. Each trial consisted of programmed acceleration from 4 to 40 RPM over a period of 3 minutes and were separated by 120 minutes. The time spent on the rotarod was recorded for each trial. Statistical significance was defined as p < 0.05 by two-way ANOVA with Tukey’s post-test and performed in GraphPad Prism v8.4.1(460) (GraphPad Software LLC., San Diego, USA).

### Inverted screen test assay

At P38, all mice underwent strength and coordination testing using the inverted screen test (Kondziella, 1964). Mice were placed in the center of a wire mesh, which was then rotated into the inverted position (mouse headfirst) and held approximately 30 cm above a padded surface for a maximum of 120 seconds. The time spent in the inverted position was recorded for each mouse. Statistical significance was defined as p < 0.05 by one-way ANOVA with Tukey’s post-test and performed in GraphPad Prism v8.4.1(460) (GraphPad Software LLC., San Diego, USA).

### Quantitative analysis

All image and quantitative analysis were performed with Olympus VS-Desktop software and ImageJ 1.52p (NIH, Bethesda, Maryland, USA) software. Sections underwent extended focal imaging (EFI) at 10X and 20X with the Olympus VS-120 slide-scanner microscope and confocal imaging at 20X and 40X. To evaluate the local effect of hemorrhage at P8, areas of red blood cell (RBC) aggregation were identified, and neighboring lobules were selected for analysis. Quantitative data was collected from comparable sections from each treatment group. At P8, measurements of EGL thickness (n = 2 sections/animal, 4 animals/treatment group), Calbindin^+^ PC density (n = 4 sections/animal, 3 animals/treatment group), GFAP^+^ Bergmann glial (BG) fiber crossings (n = 3 sections/animal, 3 animals/treatment group), and BrDU/Ki67^+^ labeling index (BrDU/Ki67^+^ cells/DAPI^+^ cells) were obtained (n = 3 sections/animal, 3 animals/treatment group). At P42, measurements of PC density, BG fiber crossings, and Parvalbumin^+^ molecular layer interneuron (MLI) density (n = 3 sections/animal, 3 animals/treatment group) were obtained from lobules VI-X. Statistical significance was defined as p < 0.05 by one-way ANOVA with Tukey’s post-test and performed in GraphPad Prism v8.4.1(460) (GraphPad Software LLC., San Diego, USA).

## Results

### Blood exposure at P6 caused significant abnormalities in adjacent regions by P8

As shown in **Figure 1B**, regions of blood exposure were readily identified and we limited our analyses to these regions in vermis and paravermis cerebellar lobules V-VII. Compared with aCSF at P8, blood exposure resulted in a significantly decreased EGL proliferation as measured by BrdU and Ki67 labeling (p = 0.0015 and p <0.0001, respectively) (**Figure 1C**,**D**). Blood exposure also significantly decreased EGL thickness compared to aCSF (p <0.0001) (**Figure 1 C**,**E**,**F**). We also saw significant loss of (Calbindin +) Purkinje cells in regions of blood exposure (**Figure 1 E, G**). GFAP+ Bergman glial fiber crossings were significantly increased following blood exposure compared to aCSF-buprenorphine (p < 0.0001) (**Figure 1E**,**H**) with erratic Bergman glial fiber branching and altered fiber architecture, compared to the typical ordered fiber structure seen in the aCSF groups. Additionally, breaks in Bergman glial fibers and endplate detachment were noted (**Figure 1E and not shown**). There were no differences between blood exposed animals that received buprenorphine vs. ketoprofen for BrdU labeling, Ki67 labeling, EGL thickness, Purkinje cell density, and Bergman glial crossings (p = 0.9729, p = 0.7645, p = 0.63, p = 0.6873, and p = 0.9531, respectively).

### Regional morphological changes were persistant at P42

Similar to findings at P8, blood exposure was associated with decreased Purkinje cell density at P42 (p < 0.0001) (**Figure 2A**,**B**). Blood exposure was also associated with decreased molecular layer interneuron density compared to aCSF (p = 0.0058) (**Figure 2A**,**C**). There was no significant difference between blood exposed animals that received ketoprofen vs. buprenorphine for PC density and MLI density (p = 0.9665 and p = 0.7783, respectively) (**Figure 2A-C**).

**Figure 2.**
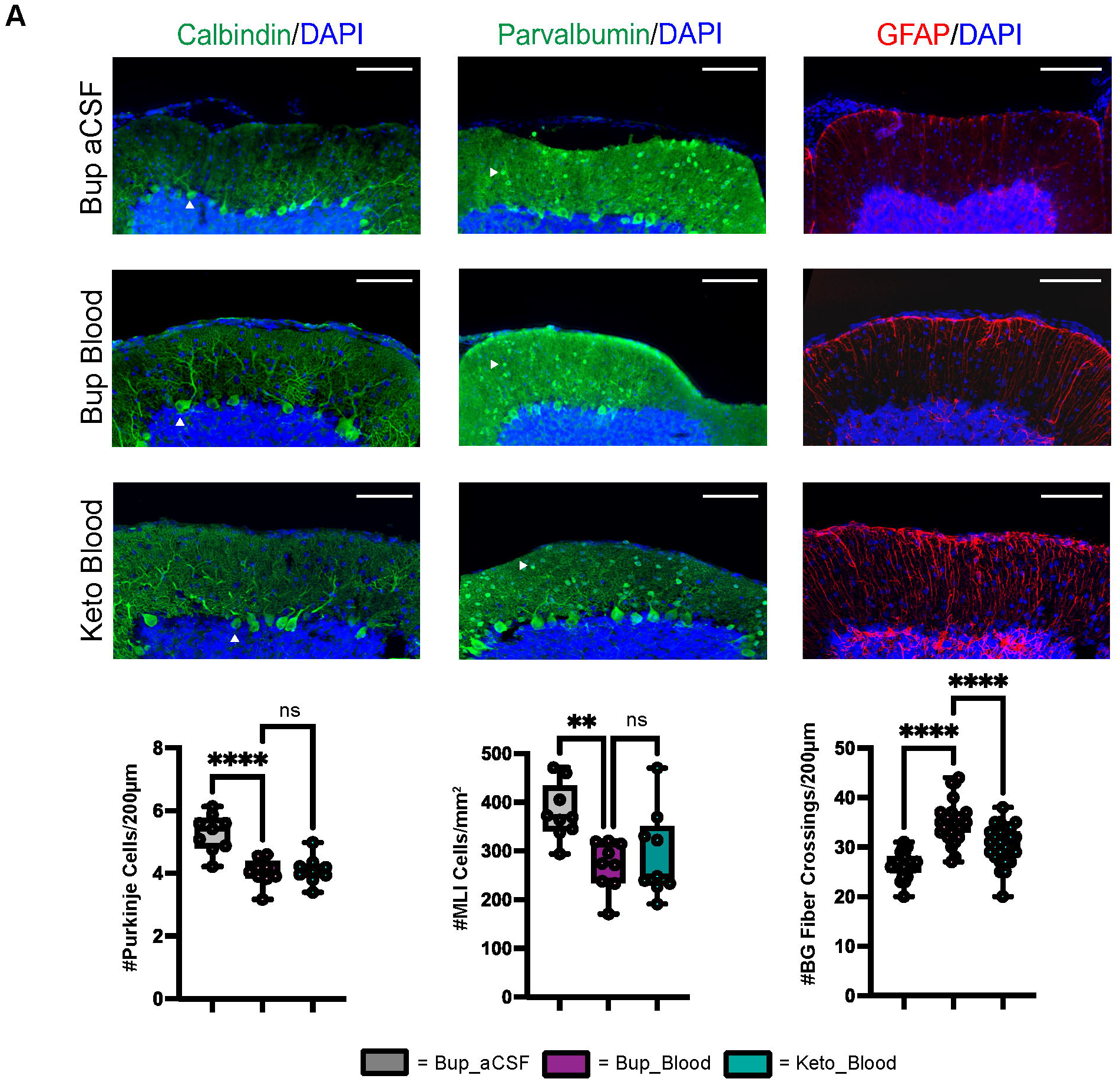
CHI results in decreased PC density, decreased MLI density, and increased BG fiber crossings within the ML at P42. **(A)** Representative confocal images at 20X demonstrating difference in PC (white arrowhead) density (left), MLI (white arrowhead) density (middle), and BG fiber density and morphology (right). Scale bars are 100 *μ*m. **(B)** Quantification of PC density in lobule VI-X for all treatment groups (N = 3, n = 9). **(C)** Quantification of MLI density in lobule VI-X for all treatment groups (N = 3, n = 9). **(D)** Quantification of BG fiber crossings within the ML for all treatment groups (N = 3, n = 27). Significance defined as p < 0.05 by one-way ANOVA with Tukey’s post-test. Data are presented as SEM with minimum and maximum values. N = number of animals per group included for analysis, n = number of section segments per group included for analysis.

Similar to findings at P8, blood exposure was associated with abnormal Bergman glial fiber branching and increased fiber crossings in animals that received buprenorphine at P42 (p<0.0001) (**Figure 2A**,**D**). Blood exposed animals that received ketoprofen had significantly fewer Bergman glial fiber crossings compared to those receiving buprenorphine (p < 0.0001) (**Figure 2D**). Blood exposure did not result in significant difference in Bergman glial soma density, either in typical position or ectopic position within the molecular layer (**Supplemental Figure 1**).

### Neonatal blood exposure did not result in significant gross motor learning/coordination deficits in older animals

Results from the inverted screen test demonstrated no significant differences between the blood and aCSF animals receiving buprenorphine (p = 0.1580); however, animals in the blood-ketoprofen group were noted to remain suspended for significantly longer than animals in the aCSF-ketoprofen group (p = 0.024) (**Figure 3B**). There were no significant differences noted between groups on Rotarod testing (**Figure 3 C**,**D**).

**Figure 3.**
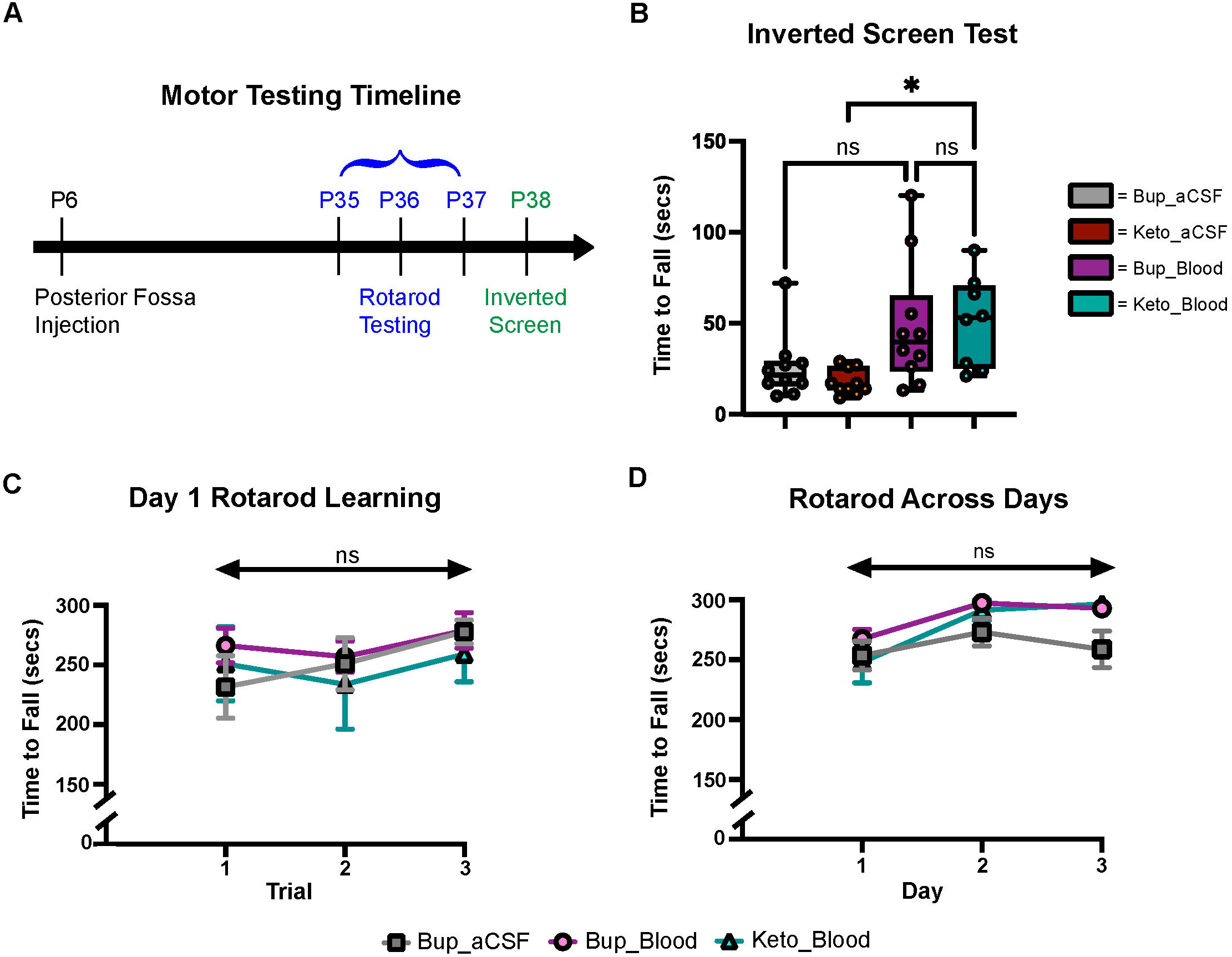
CHI did not result in significant motor learning/coordination deficits. **(A)** Schematic of motor testing assays and timeline. **(B)** Quantification of time to fall for the inverted screen test for all groups (N = 10*)^†^. **(C)** Quantification of the time to fall for Day 1 of Rotarod training for all groups (N = 10*)^‡^. **(D)** Quantification of the time to fall for Days 1-3 of Rotarod training for all groups (N = 10*)^‡^. Data are presented as mean±SEM (B) or SEM with minimum and maximum values (C,D). N = number of animals per group included for analysis. *Keto_Blood group with N = 8. ^†^ Significance defined as p < 0.05 by one-way ANOVA with Tukey’s post-test. ^‡^ Significance defined as p < 0.05 by two-way ANOVA with Tukey’s post-test.

## Discussion

To examine how preterm and neonatal cerebellar bleeds during maximal EGL proliferation in humans alters cerebellar development, we developed a novel murine model of isolated posterior fossa subarachnoid hemorrhage at P6, when maximal EGL proliferation occurs in mice. Subarachnoid blood exposure resulted in immediate and sustained morphological abnormalities to multiple cell populations immediately adjacent to the blood, especially granule and Purkinje cells, Bergman glia and molecular layer interneurons.

Multiple studies have demonstrated that Purkinje cells are exquisitely sensitive to injury, with hypoxic and ischemic insults resulting in damage to Purkinje cells in both animal models and human studies (Barenberg et al., 2001; Hausmann et al., 2007; Kántor et al., 2007; Bartschat et al., 2012; Sathyanesan et al., 2018). Importantly, Purkinje cells are essential to the normal development of the cerebellum, as Purkinje cell-derived Sonic hedgehog (Shh) protein is a key driver of granule cell precursor proliferation within the overlying EGL (Smeyne et al., 1995; Dahmane and Ruiz i Altaba, 1999; Wechsler-Reya and Scott, 1999; Lewis et al., 2004; Fleming and Chiang, 2015; De Luca et al., 2016). In the present study, subarachnoid hemorrhage was modeled at P6-P8, a key time point in cerebellar development characterized by rapid expansion of GCs in the EGL and active Shh expression by PCs. Following CHI, we found localized thinning of the EGL and decreased EGL BrDU/Ki67 labeling at P8 (**Figure 1**). We also saw immediate localized loss of Purkinje cells, with significant decreased density at P8 that was sustained at P42 (**Figures 1-2**). This result is in contrast to Yoo et al. 2014, who reported no significant change in Purkinje cell count in adult mice after induction of neonatal cerebellar hemorrhage with a bacterial collagenase injection model (Yoo et al., 2014). One caveat to our study is that we injected adult vs age matched blood, We cannot distinguish if loss is due to a direct effect on each population independently, however, loss of Purkinje cell derived Shh mitotic most certainly contributed to diminished granule cell precursor proliferation.

Recently, Bayin and colleagues demonstrated that the developing mouse cerebellum has intrinsic capacity for PC regeneration/replenishment by immature PC precursors (Bayin et al., 2018). The regeneration capacity of these cells decreases during the first postnatal week at approximately P5 (Bayin et al., 2018). In this study, blood exposure at P6 resulted in persistent decreases in PC density at P42, demonstrating significant long-term effect of CHI occurring at this developmental timepoint on the PC population (**Figure 2B**). This is important, as PC loss has been associated with multiple neurodevelopmental disorders in children, including autism spectrum disorder (ASD), many of which are present in ELGANs post cerebellar hemorrhage (Fatemi et al., 2002; Jeong et al., 2014; Skefos et al., 2014). Additionally, this finding is supportive of the limited replenishment capacity of the mouse cerebellum after the first postnatal week.

In addition to significant PC loss, our results demonstrate CHI has long-term effects on MLIs, with significantly decreased MLI density noted at P42 (**Figure 2C**). Interneuron precursor cells undergo significant proliferation and migration during development of the cerebellum, a process which is supported by PC (Fahrion et al., 2013; Galas et al., 2017). Specifically, PC-derived Shh has been demonstrated to regulate neural stem-cell like primary progenitors within the WM which give rise to inhibitory interneurons (Fleming et al., 2013). Later in development, two subtypes (Stellate cells and Basket cells) will migrate into the cortex, a process which is active in mice until P16 and may be disrupted by early postnatal insults (Zhang and Goldman, 1996). Recently, data from Sergaki and colleagues demonstrated that neurotrophic factors expressed by PCs help regulate MLI survival, suggesting that damage to PCs in the postnatal cerebellum may also result in decreased MLI survival (Sergaki et al., 2017). We postulate that CHI induced PC loss may result in impaired interactions between PCs and MLIs, resulting in diminished development of the MLI population.

BG are specialized unipolar astrocytes which have been found to guide GC precursor migration from the EGL to the IGL and may contribute to PC dendrite maturation (Rakic, 1971; Yamada et al., 2000; Lippman et al., 2008; Cheng et al., 2018). We found that CHI was associated with increased BG fiber crossings and abnormal fiber architecture at both P8 and P42 (**Figure 1E**,**H** and **2D**), but no difference in BG soma density (**Supplemental Figure 1A**). These data suggest that CHI primarily affects the branching of BG fibers rather than proliferation or maintenance of the BG population. The increase in fiber crossings likely represents reactive gliosis in response to CHI, as immature BG have previously been shown to be capable of increasing GFAP expression in response to injury (Lafarga et al., 1998).

Despite effects on multiple cell types within the developing cerebellum, we did not see statistically significant motor deficits on Rotarod or inverted screen test assays following CHI (**Figure 3B-D**). Subjectively, mice in the CHI treatment groups were noted to spend more time clinging to the rotating rod and less time demonstrating coordinated walking during the Rotarod trials compared to the aCSF mice. Additionally, mice in the aCSF groups exhibited increased exploratory behavior during the inverted screen test, while CHI mice tended to remain in a small area with less movement. It is possible that this increase in exploratory behavior resulted in increased likelihood of grip loss and falling from the wire mesh, explaining why mice in the Keto-aCSF group fell off of the mesh more quickly than mice in the Keto-CHI group (**Figure 3B**). Moreover, it is possible that subtle motor deficits may have been found utilizing different behavior assays, such as eyeblink conditioning or vestibulo-ocular reflex analysis. This is a limitation of our current study and requires further investigation to understand the relationship between CHI and motor deficits. There is ample evidence from human cerebellar bleed patients that severity of functional deficits is dependent on injury size and topography (Brossard-Racine and Limperopoulos, 2021b).

Secondarily, we compared the use of the opiate buprenorphine and the nonsteroidal anti-inflammatory drug (NSAID) ketoprofen in order to investigate the possible role of neuroinflammation in injury. The use of ketoprofen in our model was associated with variable results, but did not demonstrate a significant neuroprotective effect. In the acute injury phase (P6-8), we found no significant difference post CHI in EGL thickness, PC density, BrDU/Ki67 labeling indices, and BG crossings (**Figure 1**) between buprenorphine and ketoprofen cohorts. Similarly, we saw no significant difference in PC density, MLI density, oligodendrocyte density, or BG soma density at P42 (**Figure 2, Supplemental Figure 1**). We did see a significant decrease in BG fiber crossings at P42 in CHI mice treated with ketoprofen compared to those treated with buprenorphine (**Figure 2D**). Taken together, these data suggest that anti-inflammatory treatment does not have a profound protective effect on cerebellar development in our model of CHI; however, additional studies are required to further elucidate the role of neuroinflammation, particularly with regards to reactive Bergmann gliosis.

In summary, we developed a novel neonatal mouse model of isolated posterior fossa SAH to model neonatal CHI, directly examining the effects of blood exposure on cerebellar development. Our data demonstrates that exposure of the developing cerebellum to blood results in immediate disruptions to multiple aspects of cellular development and most strongly suggests the involvement of PC loss in the pathophysiology.

## Author Contributions

This study was conceived by KJM, DFB, JS and CMT. All experiments were designed and performed by DFB, CMT and JS. KJM and DFB wrote the manuscript with input from all authors.

## Funding/Acknowledgements

This work was funded by NIH R37NS095733 to KJM.

**Supplemental Figure 1. CHI did not result in significant differences in BG soma density or ectopic positioning of BG soma at P42**.

**(A)** Quantification of BG soma density for all treatment groups (N = 1, n = 9).

**(B)** Quantification of BG soma density noted in ectopic positioning for all treatment groups (N = 1, n = 9).

Significance defined as p < 0.05 by one-way ANOVA with Tukey’s post-test. Data are presented as SEM with minimum and maximum values.

N = number of animals per group included for analysis, n = number of sections per group included for analysis.

