## Supplementary figures and images for "Neonatal Subarachnoid Hemorrhage Disrupts Multiple Aspects of Cerebellar Development"

### Supplemental Figure 1

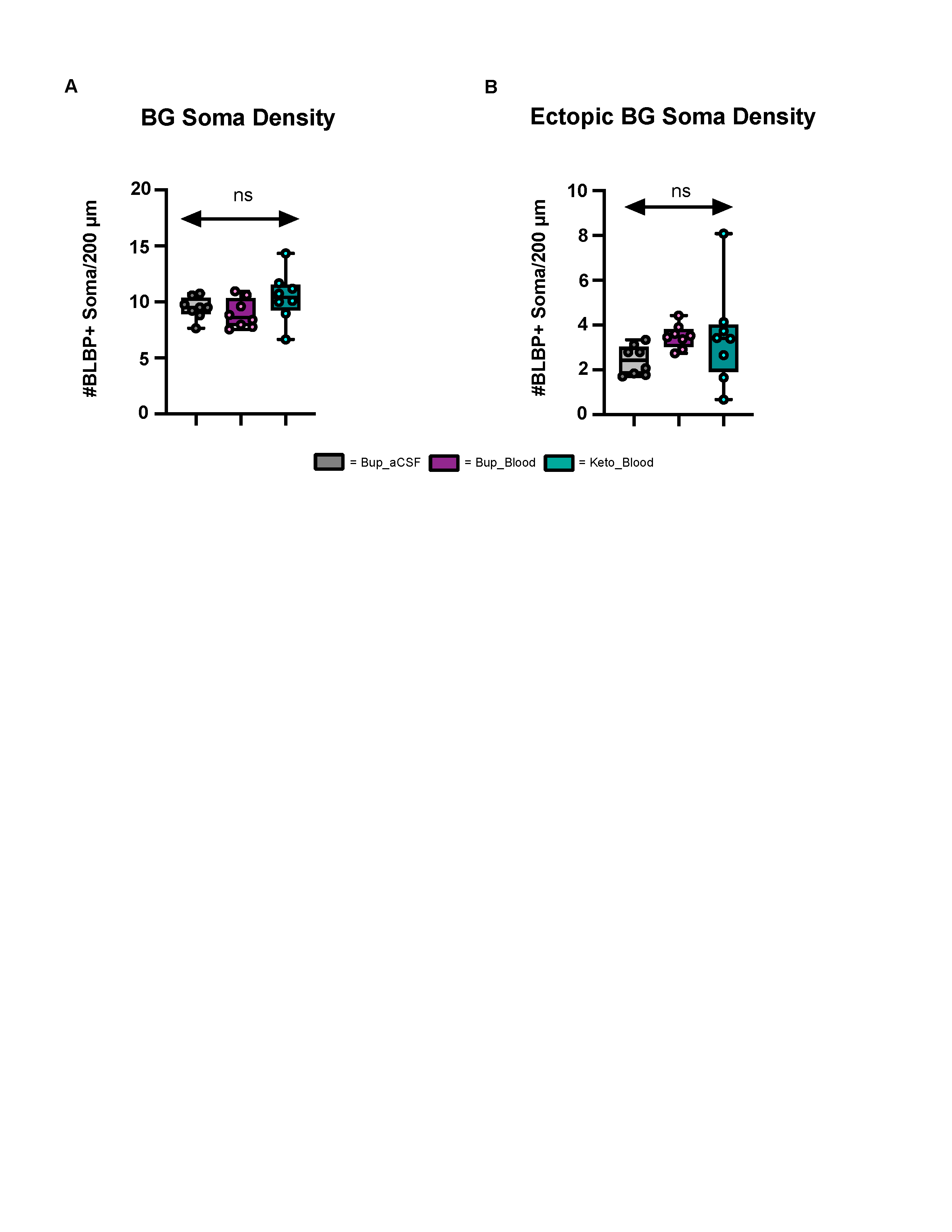
